# Reimagined Microphone-Free Acoustic Volumetry: An Open, DIY Platform for Global Phenotyping

**DOI:** 10.1101/2025.04.25.650573

**Authors:** Abbas Haghshenas, Yahya Emam

## Abstract

We present a groundbreaking Do-It-Yourself (DIY) acoustic volumetry platform that redefines quantitative measurement by eliminating the conventional microphone. Our design exploits the inherent acoustic–electrical properties of a dynamic microphone cartridge mounted on a sealed chamber—representing the theoretically simplest possible acoustic volumeter. By focusing on resonance peak shifts, our streamlined sensor —utilizing an easily assembled circuit built solely from off-the-shelf audio connectors to split the sound card output between sensor excitation and response recording—delivers rapid, precise volumetric measurements with as few as three frequency points in several seconds.

Calibration using both linear and logarithmic models demonstrated a robust correlation between resonance peak shifts and sample volume, yielding root mean square errors (RMSE) of 1.980 μL and 1.662 μL, respectively. Notably, when applied to a ten-grain assay, these values correspond to an average error of less than 0.2 μL per wheat grain, thereby affirming the device’s precision across a diverse range of sample volumes. An exclusive Python-based freeware, distributed globally, provides an intuitive interface for calibration and measurement, ensuring that this cost-effective and modular approach is accessible to researchers worldwide. This innovative method not only simplifies traditional volumetry techniques but also paves the way for further optimization, marking a significant advancement for applications across a broad spectrum of scientific fields.

## Introduction

Object volumetry has long been a cornerstone of quantitative measurement in science. Its evolution spans from classical direct and displacement methods—rooted in Archimedes’ principle (e.g., fluid displacement [Huxley, 1971; Bontzos et al., 2019], hydrostatic weighing [Hughes, 2005; Zůda, 2022], and gas displacement techniques [Tamari, 2004])—to contemporary methods such as integration techniques, stereological approaches, numerical approximations (e.g., Mattfeldt, 1987; Black, 1999), imaging-based digital volumetry (Hughes et al., 2017; Musse and Van, 2018; Xiong et al., 2019), and acoustic volumetry (Lao, 1978; Kobata et al., 2004; Sydoruk et al., 2020).

Acoustic volumetry, in particular, offers several distinct advantages. It is simple, inexpensive, accurate, fast, and noninvasive since it does not require immersing the object. Most acoustic volumeters reported to date comprise three primary components: (1) a closed container (which may be either simple or complex), (2) an excitation unit (typically a speaker that propagates sound waves within the container), and (3) a detection unit (one or more microphones that capture the propagated sound waves) (e.g., Kobata et al., 2004; Sydoruk et al., 2020). In these devices, irrespective of the relative positions of the speaker, microphone, and object—whether the object is placed between the speaker and the microphone (as described by Kobata et al., 2004) or the speaker is located between the object and the microphone (as reported by Sydoruk et al., 2020)—the fundamental measurement mechanism remains unchanged. When an object is placed in the measurement chamber, its effective volume decreases; driving the speaker at the same level therefore produces smaller pressure oscillations, which reduce the electromotive force in the microphone coil and result in a lower sound amplitude recorded by the microphone than in the empty-chamber reference. In this context, Sydoruk et al. (2020) introduced a notable sensor configuration in which the speaker is positioned in-line between the object and the microphone within a cylindrical chamber. This design enabled a detailed analysis of the force interactions among the sensor elements and the surrounding air, as well as the force losses attributable to the membranes of both the microphone and the speaker. It is anticipated that simplifying the sensor’s architecture—by reducing the number of intermediate cavities between the object and the membranes, or even by eliminating one of the membranes—could mitigate unwanted factors such as resistances and energy loss, thereby potentially enhancing the accuracy of volume estimation.

Conversely, the simplest acoustic volumeter reported is also the oldest design (see Lao, 1978), which involves installing a single transducer on a sealed container. In this configuration, the acoustic transducer is integrated into a dedicated circuit that monitors voltage variations resulting from changes in impedance within the container as it transitions from an empty to a nonempty state. However, according to both the literature and market evidence, this type of volumeter has not been widely utilized in the various scientific disciplines that require volumetric analysis over the past half-century.

In this study, we propose the development of a novel, reliable, and DIY acoustic volumeter designed for global use. By removing the microphone, we seek to deliver a reimagined version of the simplest acoustic volumeter directly to researchers in diverse fields. Our approach refines the estimation methodology and incorporates user-friendly software, with particular emphasis on applications in crop physiology and phenotyping.

## Materials and Methods

### Sensor

The acoustic volumeter sensor featured a simple design (Fig. 1). An inexpensive dynamic microphone cartridge (EOEL, manufactured in China) was affixed using silicone glue to an annular cylindrical plexiglass container. The container has an outer diameter of 70 mm, a central hole of 25 mm in diameter (matching the cartridge’s effective diameter), and a thickness (height) of 10 mm. The cartridge was connected to a 3.5 mm audio jack with high-quality wiring. Placing the sensor on a flat, polished surface, such as scratch-free glass, formed a temporary sealed chamber into which a sample could be placed for volumetry.

**Figure 1.**
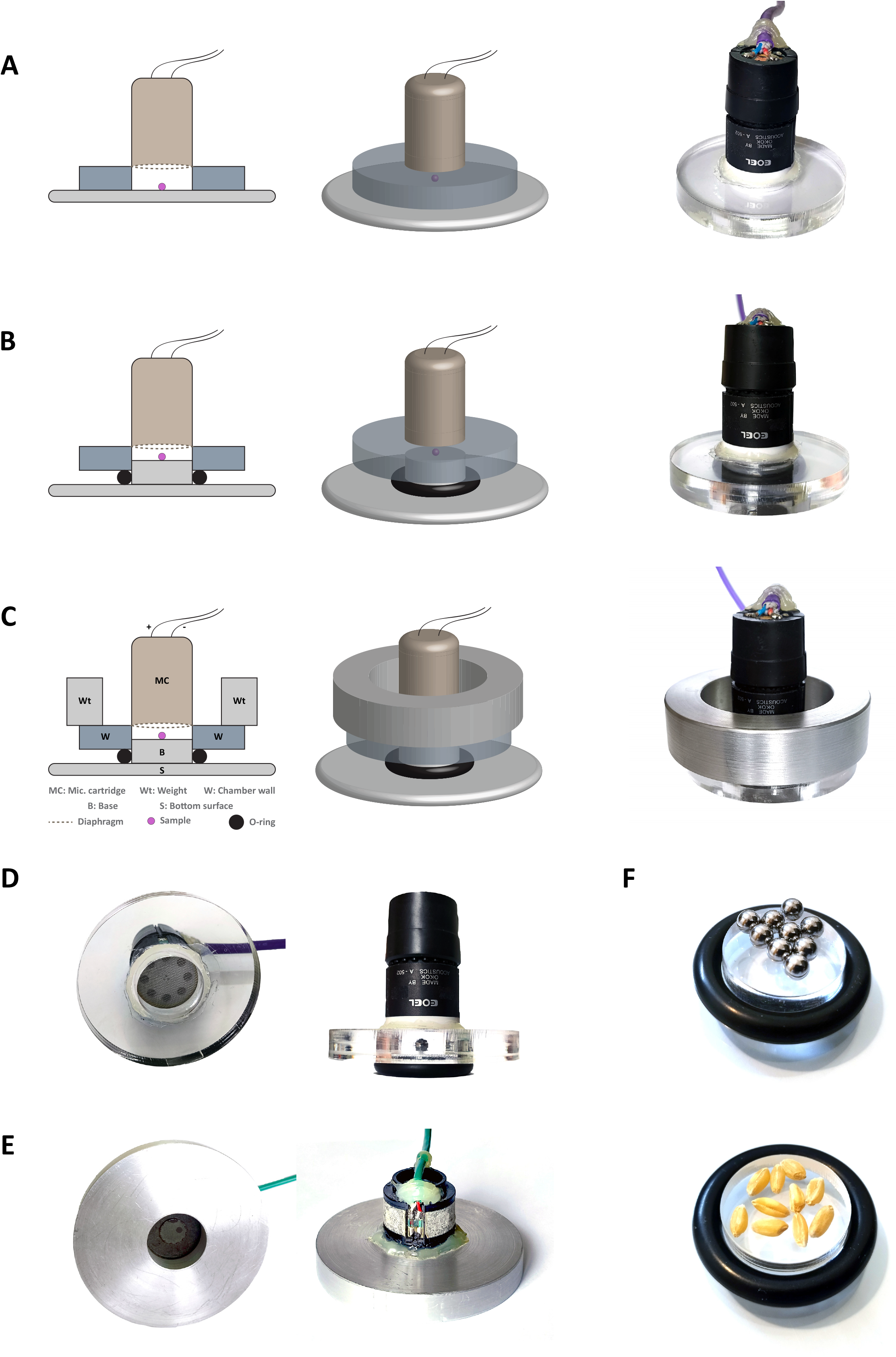
Structure of the Microphone-Free Acoustic Volumeter Sensor. (A) Basic sensor design comprising an acoustic diaphragm (e.g., a microphone cartridge) mounted on a chamber. (B) Improved sensor design featuring a sealing O-ring and an additional base to enhance accuracy by reinforcing the chamber seal. (C) Final, most advanced sensor design, incorporating an O-ring, additional base, and added weight for optimal sealing and accuracy. (D) Alternative viewing angles of the sensor. (E) A variant of the microphone-free acoustic volumeter constructed from alternative materials (e.g., aluminum) with a different cartridge. (F) Calibration and measurement setups, showing 5 mm steel balls arranged on the plexiglass base (for sensor calibration) and wheat grains placed on the same base (as volume measurement samples).

Extensive pre-analyses were performed to optimize the sensor chamber’s sealing. Various techniques—including applying a few droplets of distilled water between the surfaces and using handmade silicone washers—were tested (data not shown). Ultimately, the best sealing was achieved by using a donut-shaped rubber O-ring (37 mm outer diameter, 25 mm inner diameter, 6 mm thick) combined with an additional weight of 160 g, bringing the total sensor weight to 230 g. This configuration maintained a consistent and reliable acoustic response (see Fig. 1, parts B and C). Additionally, a cylindrical plexiglass base (25 mm diameter, 8 mm height) was employed to secure the washer on the bottom surface and to reduce the sensor chamber’s empty volume. Alternative sensor versions using different materials and cartridge brands were also constructed and evaluated (see Fig. 1, parts D and E); however, the results presented herein pertain exclusively to the sensor described above, unless otherwise specified.

### Circuit arrangement

A standard onboard PC sound card (Realtek® Audio) with driver RTKBHD64.sys (version 6.0.8703.1, Realtek Semiconductor Corp., May 14, 2019) was used for both generating acoustic signals and recording the sensor’s response. The default sound format was configured to 16-bit depth at a 44.1 kHz sampling rate via the Realtek control-panel settings. Figure 2 illustrates the direct connection of the sensor to the sound card’s input and output ports. The output from the sound card’s headphone jack was split into two paths: one driving the sensor and the other routed back to the sound card via the microphone input. This configuration effectively connects the sensor in parallel with the microphone input. The experiments were conducted on a computer running Windows 10 Pro 64-bit (version 10.0, Build 19041, Microsoft) built around a Gigabyte Technology Co., Ltd. B250-HD3P-CF motherboard. Before each experiment, microphone playback was muted in both the Windows sound settings and the Realtek control panel.

**Figure 2.**
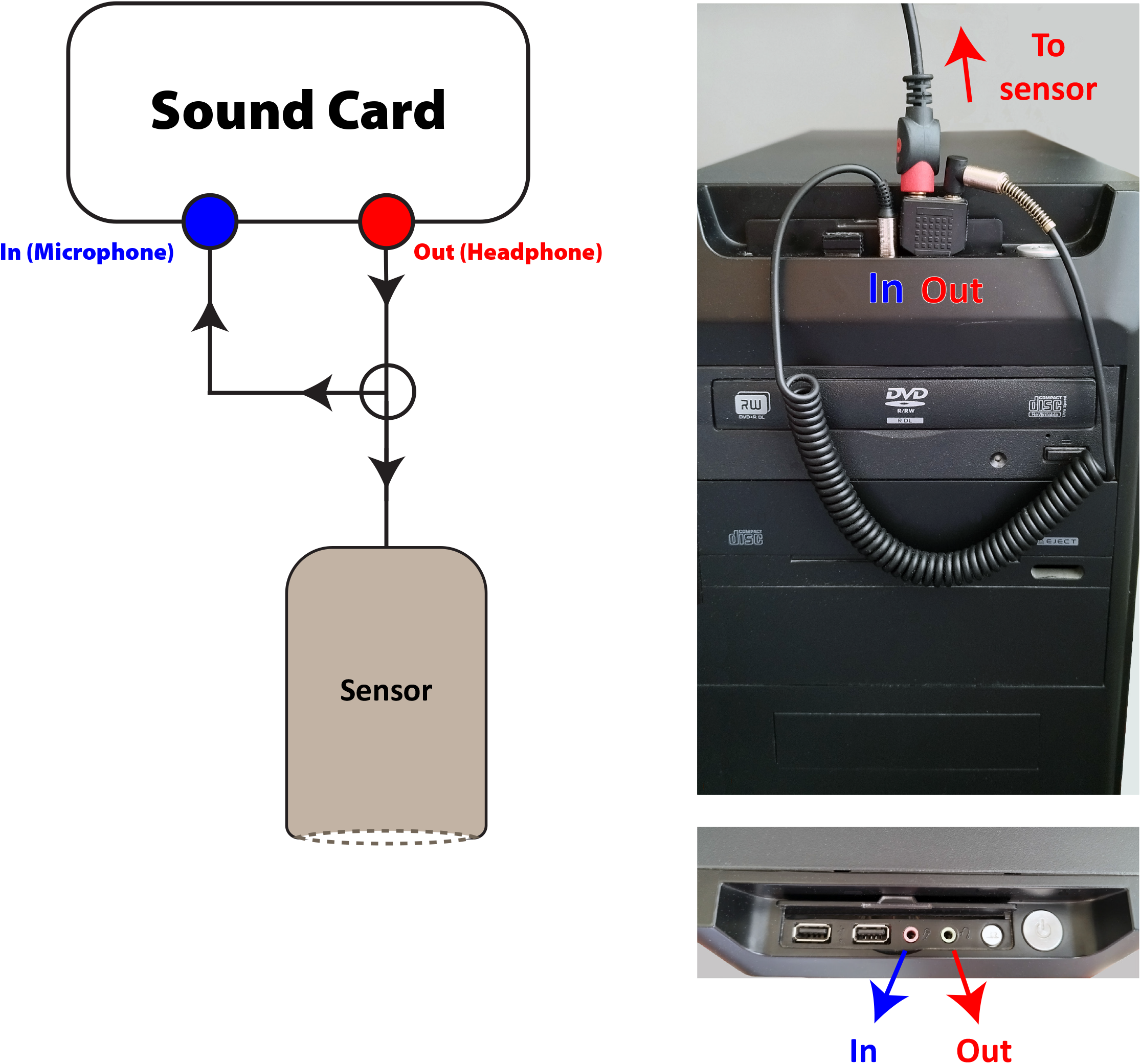
Connection configuration of the acoustic volumeter sensor to the PC sound card. The output from the headphone jack is split into two paths: one leads to the sensor, while the other returns to the sound card via the microphone jack. This arrangement establishes a parallel connection between the sensor and the microphone input.

### Computation

Preliminary analyses were performed using Audacity 3.6.4. All subsequent advanced evaluations— including acoustic profiling, calibration, modeling, and volumetry—were conducted with a custom-developed, Python-based freeware named Acoustic Volumeter v.1 (available at https://github.com/haqueshenas/Acoustic-Volumeter).

#### Acoustic Profiling

To determine the sensor’s acoustic response, an empty chamber was subjected to a series of sinusoidal sound frequencies generated using the Profiling tool of the Acoustic Volumeter software. The following settings and specifications were applied:

- Frequency range (Hz): Typically from 100 to 3000 Hz (adjustable as needed)
- Playback level (0–1): Amplitude of the generated sound
- Playback duration (s): Duration of each frequency’s playback and recording
- Frequency resolution (Hz): Interval between consecutive frequencies
- Silence duration (s): Interval of silence between successive frequency playbacks
- An option to exclude the initial milliseconds (latency) from the root mean square (RMS) amplitude calculation for each frequency.

#### Sensor Calibration

The sensor calibration was conducted in two steps (Fig. 6). In the first step, the resonance peak of the sensor was determined under various conditions, ranging from an empty chamber to conditions with one or multiple samples (i.e., steel balls of 5 mm diameter). Simple models were fitted to the distinct convexity observed in the profiling graph. The following models were evaluated:

##### Quadratic

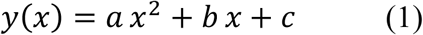

where *a, b*, and *c* are the polynomial coefficients.

##### Gaussian

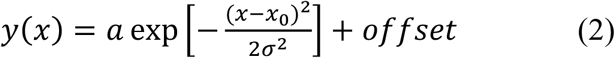

###### Parameters

- *a*: Peak amplitude (initialized as max(*y*))
- *x*_0_: Center (initialized as mean(*x*))
- σ: Width (initialized as std(*x*))
- *offset*: Baseline level (initialized as min(*y*))

##### Lorentzian

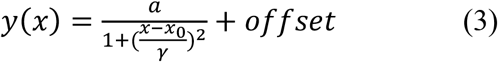

###### Parameters

- *a*: Amplitude
- *x*_0_: Center
- γ: Related to the half-width (initialized as std(*x*))
- *offset*: Baseline (initialized as min(*y*))

##### Asymmetric Lorentzian

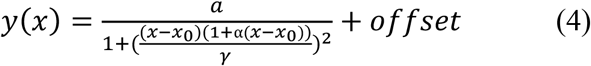

###### Parameters

- *a*: Amplitude
- *x*_0_: Center
- γ: Width parameter
- *offset*: Baseline (initialized as min(*y*))
- α: Asymmetry parameter (initialized as 0.1)

##### Voigt

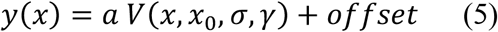

where

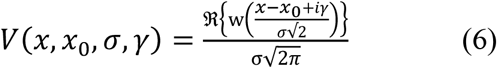

and w(z) is the Faddeeva (complex error) function.

###### Parameters

- *a*: Amplitude
- *x*_0_: Center
- σ: Gaussian width (initialized as (max(*x*) − min(*x*))/4)
- γ: Lorentzian width (initialized as (max(*x*) − min(*x*))/8)
- *offset*: Baseline (initialized as min(*y*)

In the second calibration step (Fig. 6), the set of resonance peaks, collected under conditions ranging from an empty sensor chamber to a chamber containing up to nine sample balls, was used to model the overall trend in resonance peak variation. Various trend functions were evaluated, including linear, simple quadratic, cubic, and logarithmic fits, as well as a piecewise linear model (hereafter referred to as “Seg-linear” i.e. segment linear). In the Seg-linear model, independent linear equations were fitted between each pair of consecutive calibration points. During measurement, each data point was assigned to its corresponding segment, and the appropriate linear function was used to estimate its value, enabling localized linear approximation across the entire dataset.

Once calibration was complete, the calibration outputs were used for volumetric measurements, including validation tests and volumetry of wheat grains using the Measurement tool of the Acoustic Volumeter software. All statistical analyses were conducted using XLSTAT (Version 2016.02.28451; Addinsoft). Notably, all reported procedures—including sensor acoustic profiling, resonance peak modeling, overall trend calibration, and measurement—are fully automated and user-friendly within Acoustic Volumeter v.1. All evaluations were carried out at an ambient temperature between 17– 19 °C.

## Results

Figure 3 shows that the recorded signal is in anti-phase relative to the original output waveform, confirming the sensor structure’s bidirectional acoustic-electrical behavior. Further evidence of the sensor design and circuit configuration’s ability to capture the chamber’s acoustic properties is provided in Figure 4. In this figure, regardless of the transducer’s material, model, size, or recording specifications, various sensor versions consistently demonstrated a distinct resonance convexity at approximately 1000 Hz (noting that the exact frequency may vary with the transducer model or sensor characteristics).

**Figure 3.**
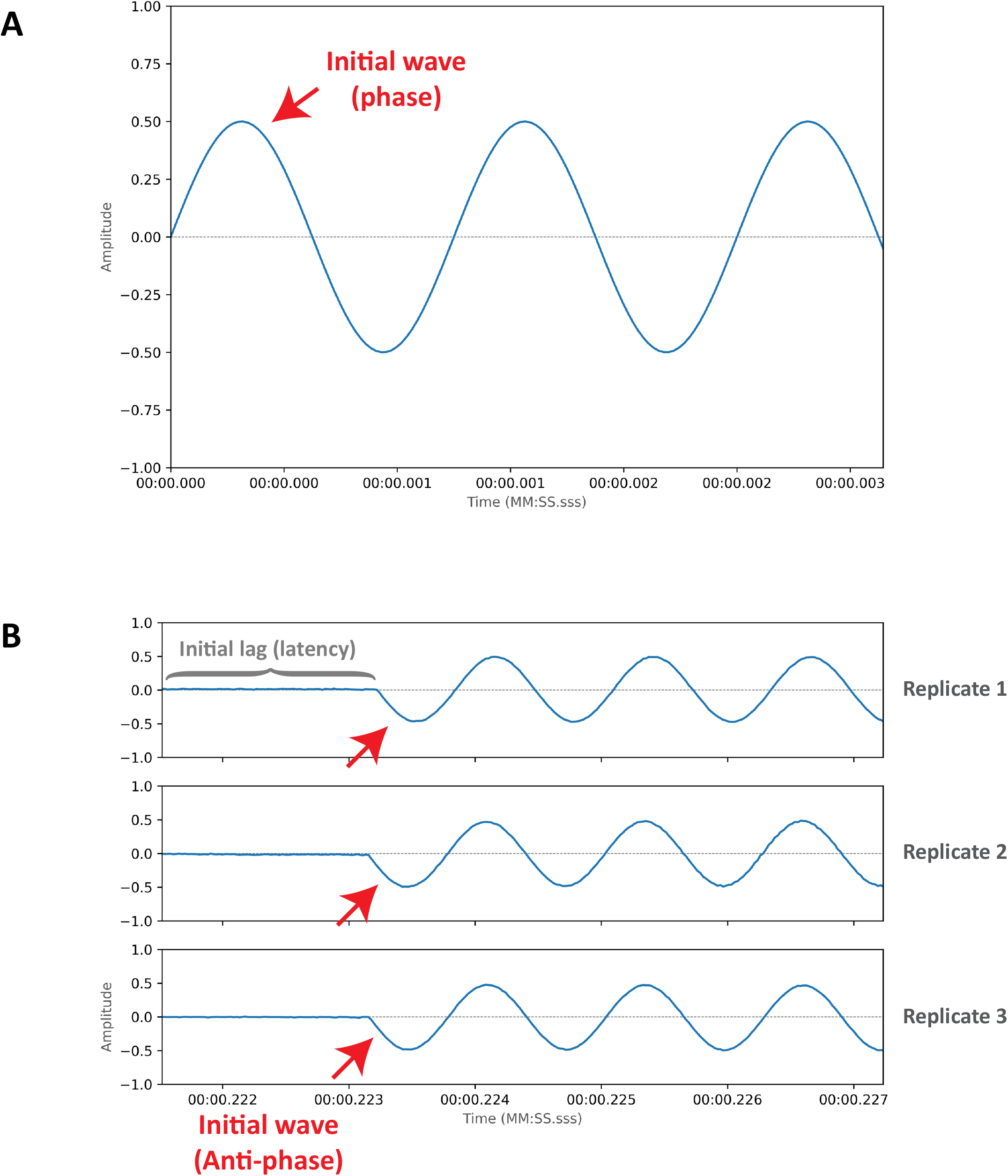
Comparative representations of sinusoidal waveforms (800 Hz, 50% amplitude) for (A) the played (in-phase) sound and (B) the recorded (anti-phase) sound in the microphone-free volumeter. Only three replicates of the recorded signal are shown; however, all recordings made using this sensor exhibit the same waveform, starting with negative amplitude values as an anti-phase response. This behavior is analogous to that observed in systems with a conventional microphone, where the diaphragm is initially displaced backward.

**Figure 4.**
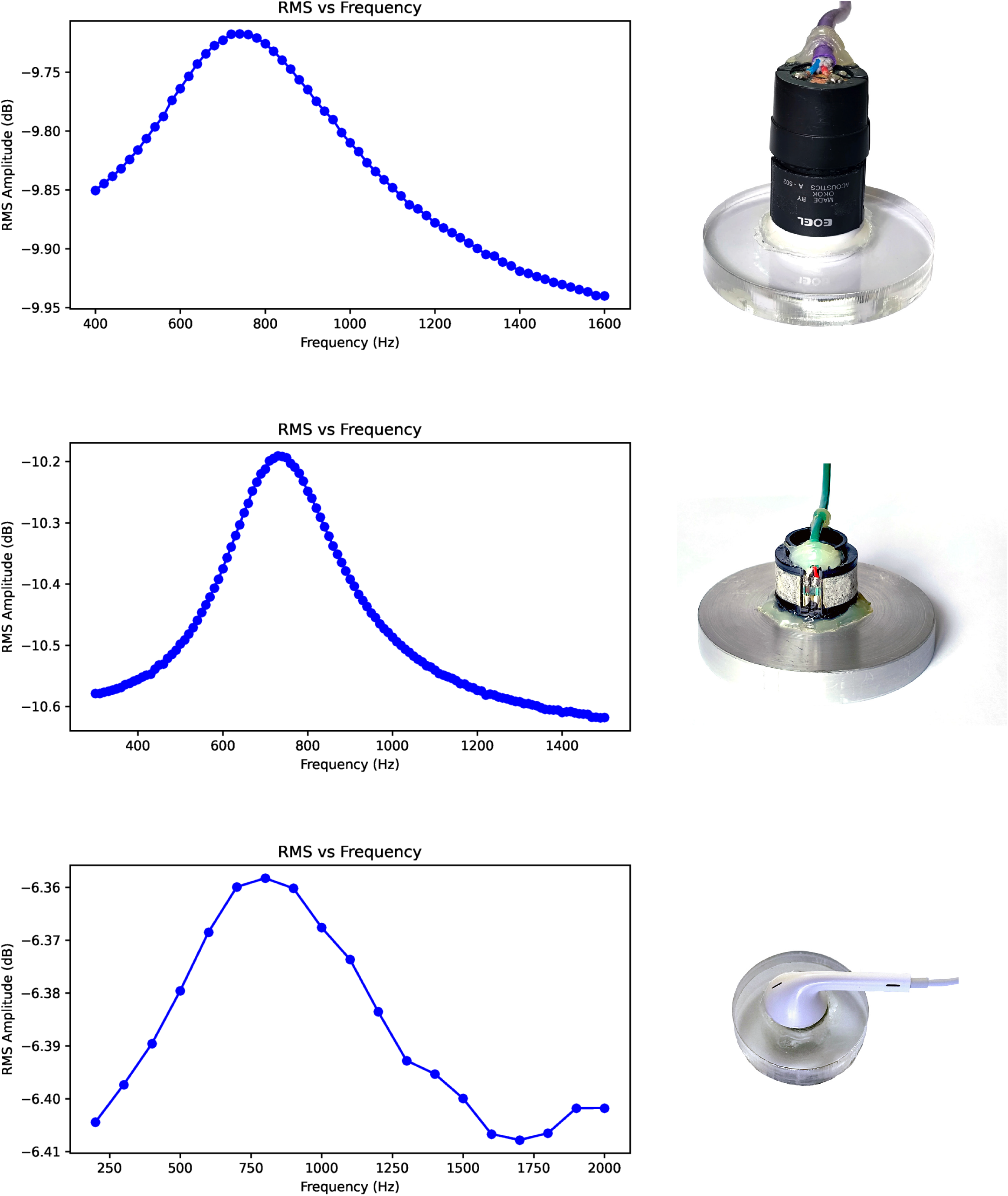
Acoustic profiles of three different sensors equipped with various EOEL and Gilson microphone cartridges and an Apple earbud, respectively. In the recorded graphs of all sensors, a single distinguishable peak (i.e., the resonance frequency) is clearly detectable below 1000 Hz. Notably, the sound was played and recorded for each sensor according to its specific design, and each data point represents the result of a 5-second recording of a single tone.

As an additional validation of the sensor’s capability, Figure 5 illustrates that increasing the sample volume (thereby reducing the internal chamber volume) resulted in a shift of the resonance convexity and its peak towards higher frequencies. This behavior confirms the sensor’s sensitivity to volumetric changes.

**Figure 5.**
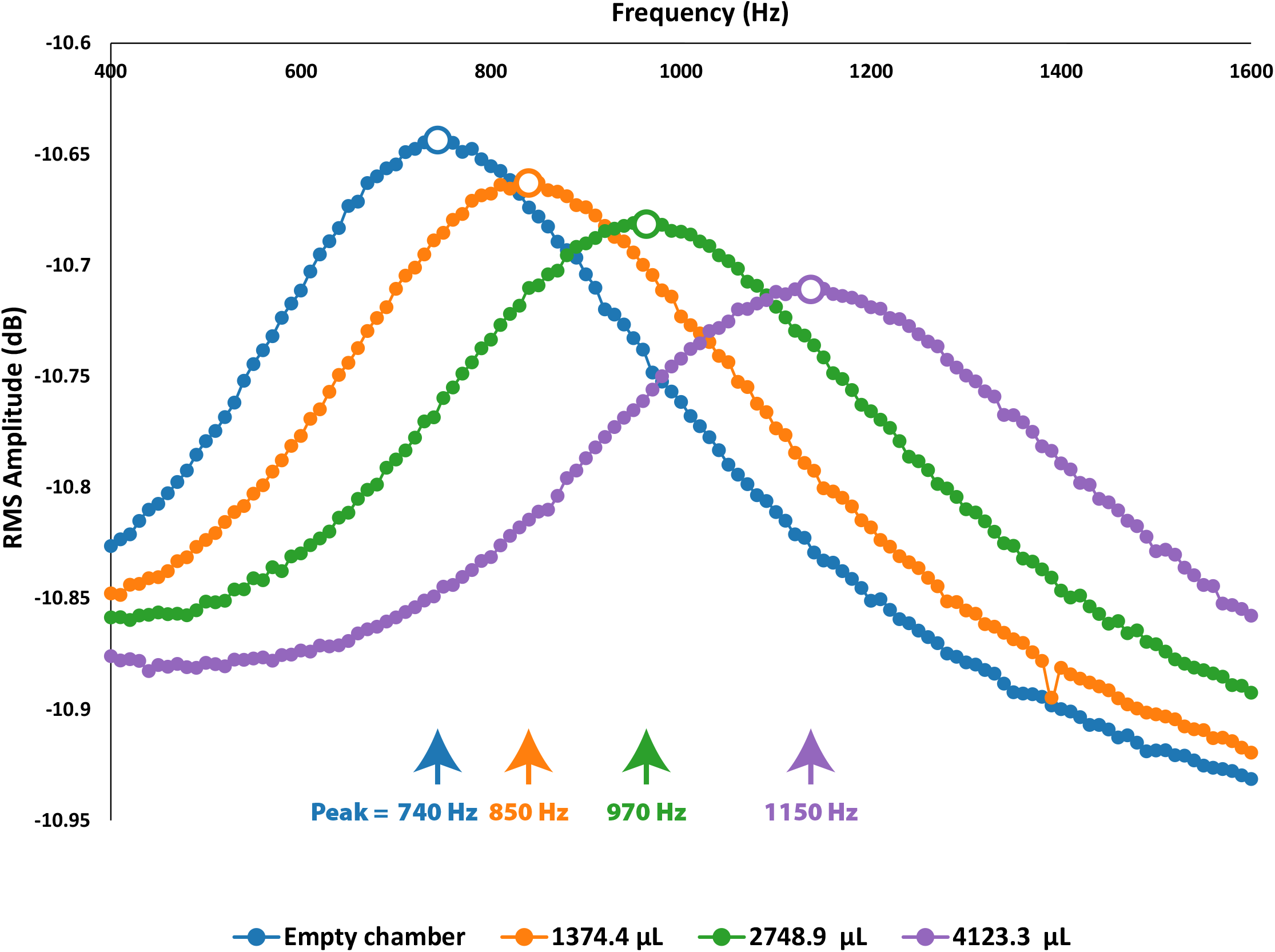
Recorded resonance peaks shi to higher frequencies with increasing sample volume. The blue, orange, green, and purple curves represent the conditions in which no object, one cylindrical plexiglass disk (geometrically calculated at 1374.4 μL), two disks (2748.9 μL), and three disks (4123.3 μL) were placed in the volumeter chamber, respectively. Sampling specifications were as follows: Frequency Range, 400–1600 Hz; Playback Level, 0.01; Frequency Resolution, 10 Hz; Playback Duration, 5 s (each point represents a 5-second tone playback and recording); Silence Duration, 0.5 s; and Lag Duration, 50.0 ms.

During calibration, the resonance peaks obtained under different conditions —by introducing varying numbers of calibration balls (5 mm diameter) into the sensor chamber— were precisely estimated using different models fitted to the corresponding convexities (see Figures 6A and 6B). The overall trend of changes in these estimated peaks exhibited a robust linear relationship (R^2^ = 0.998; Fig. 6C), enabling straightforward estimation of the sample volume.

**Figure 6.**
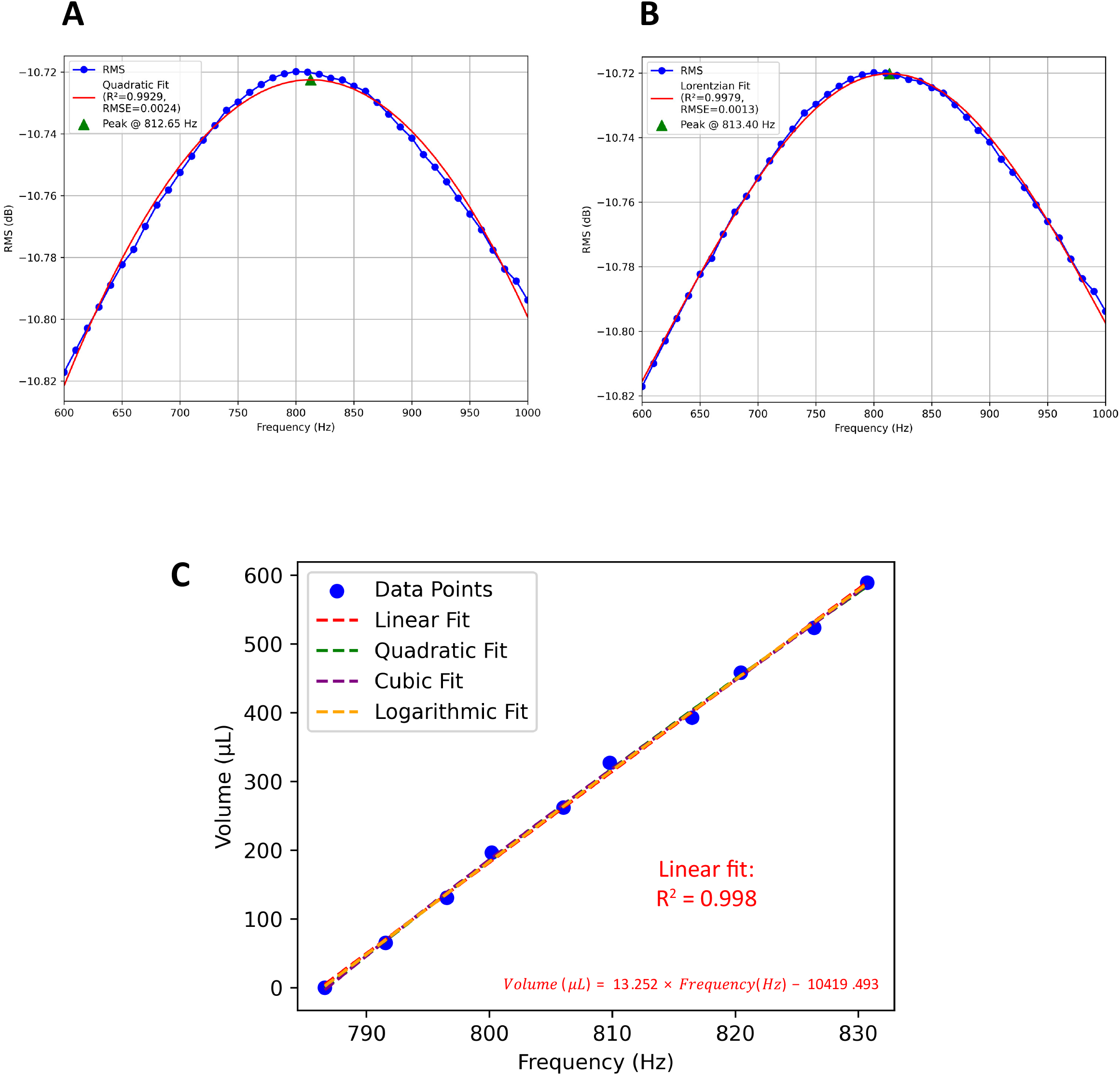
Sensor Calibration. Acoustic Volumeter v.1 freeware outputs illustrate two primary calibration steps. In the first step (A and B), the sensor’s resonance peaks are determined for various standard objects of different volumes placed in the chamber by fitting selected models (e.g., Quadratic, Gaussian, Lorentzian, Asymmetric-Lorentzian, and Voigt); here, the results of the Quadratic (A) and Lorentzian (B) fits are shown. In the second step (C), the overall trend in the shifts of the determined resonance peaks is modeled against the geometrically calculated volumes of the calibration objects. Notably, the calibration objects ranged from an empty chamber (zero volume) to nine standard steel balls, each with a 5 mm diameter. This model, which is typically linear and remains relatively consistent over a wide range of ambient conditions, is used to directly estimate object volumes based on the resonance frequency.

Figure 7 presents the validation results of the volumetry method based on shifts in the resonance peak. Among the various models tested on the overall trend, both the linear and logarithmic fits provided the most accurate estimations, with root mean square error (RMSE) values of 1.980 and 1.662 μL, respectively.

**Figure 7.**
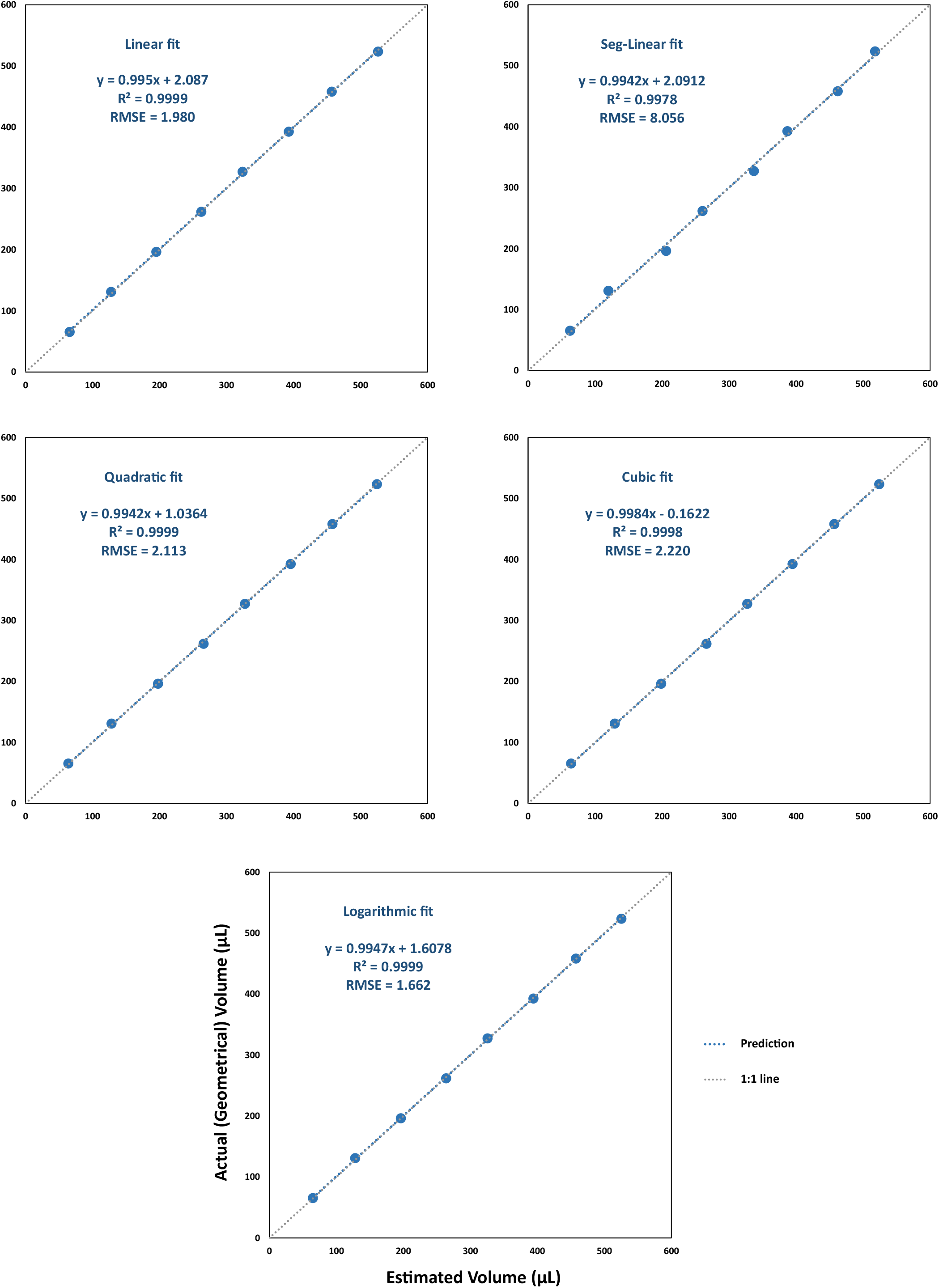
Validation of resonance-based acoustic volumetry using Linear, Segmented-Linear, Quadratic, Cubic, and Logarithmic models. Each data point represents the mean of three replicates of steel ball volumetry, obtained under the following specifications: Frequency Range, 700–900 Hz; Playback Level, 0.01; Frequency Resolution, 50 Hz; Playback Duration, 10 s; and Silence Duration, 0 s.

To explore the potential for enhanced applicability and reduced measurement time, additional analyses were performed using varying measurement specifications. The results indicated that the number of frequency points required for reliably detecting the resonance convexity and estimating its peak could be reduced to the theoretical minimum of three, even when using a short recording duration of 5 seconds per frequency (reducing the total measurement time to 15 seconds; see Fig. 8). Additionally, as depicted in Figures 9A and 9B, variability in the source signal appears to be a significant factor limiting precision in acoustic volumetry. This negative effect can be mitigated by increasing the measurement duration, as illustrated in Figure 9C.

**Figure 8.**
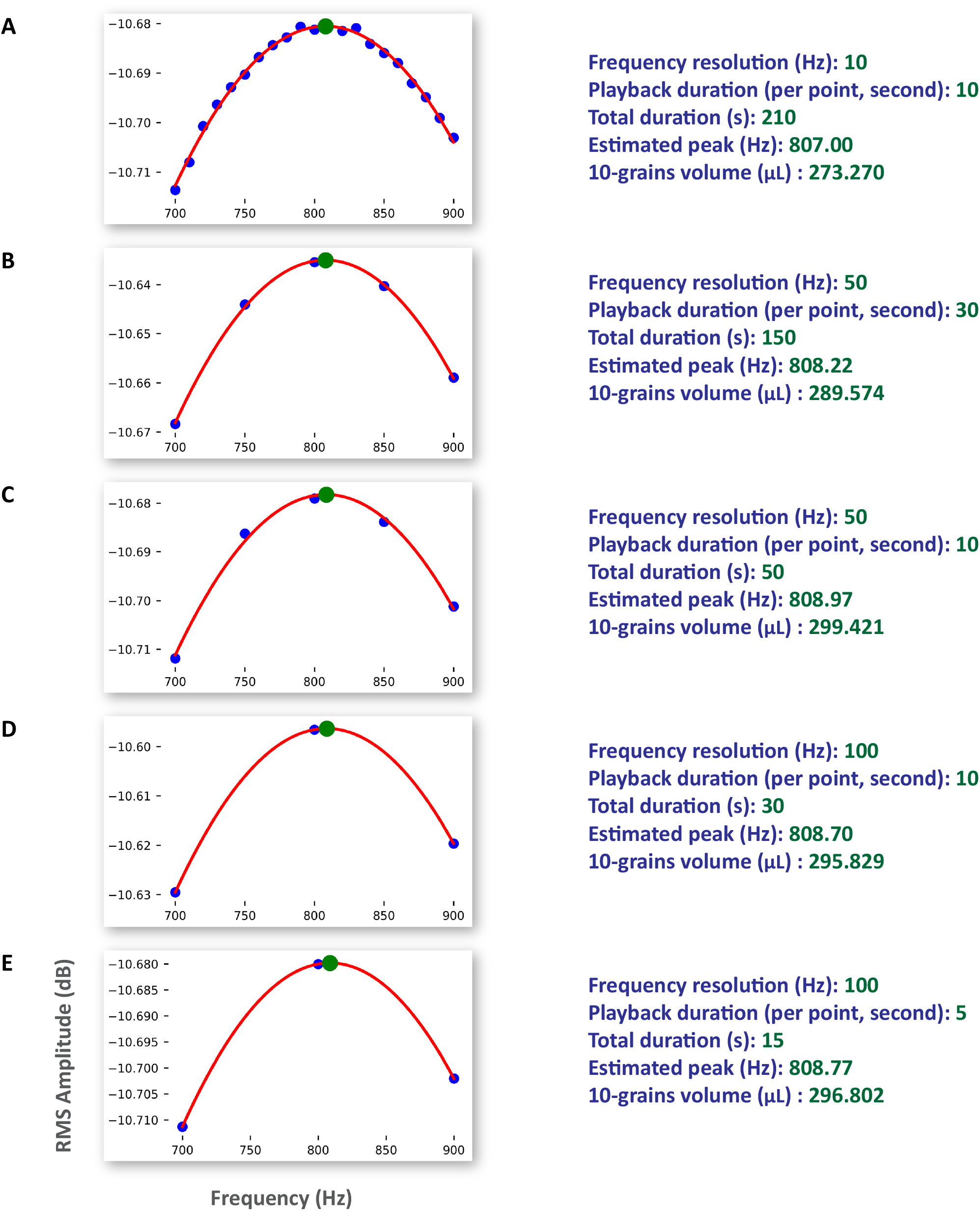
Acoustic volumetry of a 10-grain wheat sample under varying measurement specifications. Different frequency resolutions (i.e., the number of sampling points for resonance peak determination) and playback durations produced varying total measurement times, with the silence duration set to zero throughout. Resonance peaks were estimated using quadratic fits, and a single linear model was applied for volume estimation. To facilitate comparability across all sections, this linear model was calibrated using a single specification set (10 Hz resolution and 10 s playback), although volume estimation should ideally use models developed for each measurement mode. Images and results are from the outputs of Acoustic Volumetry v.1.

**Figure 9.**
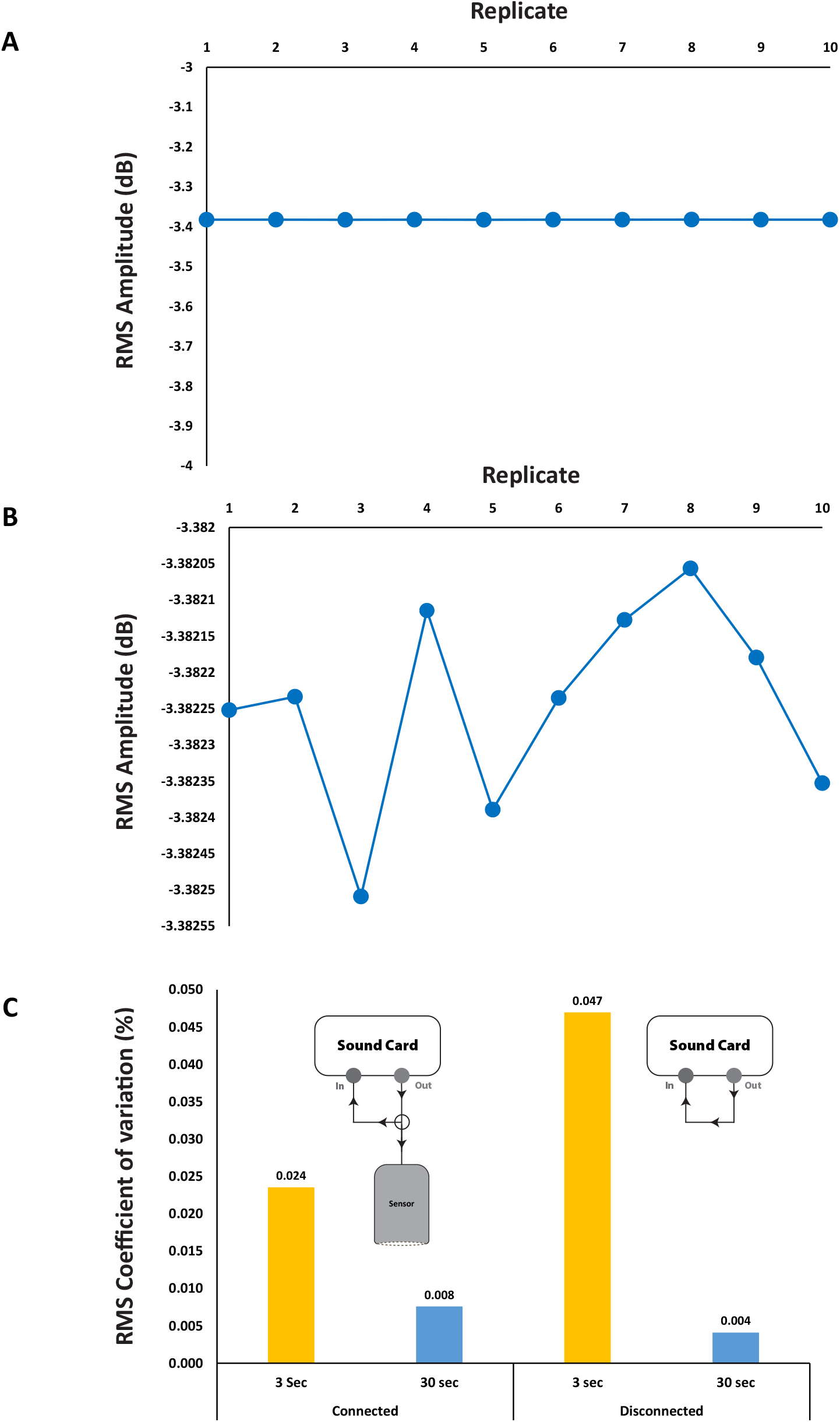
Source signal variability as a precision-limiting factor in microphone-free acoustic volumetry. Panels A and B present comparative plots illustrating the input signal level (sensor input or sound card output) for a 630 Hz tone played and recorded over 30 seconds at a normalized amplitude of 0.01 (range: 0–1) across 10 replicates, under two numerical precision settings (reflected in the vertical axis values). These measurements were conducted with the sensor disconnected from the circuit. Panel C displays the coefficient of variation (%) in the root mean square (RMS) amplitude under comparable conditions (630 Hz tone at 0.01 playback level), comparing recording durations of 3 and 30 seconds, both with the empty sensor connected and disconnected. In both cases, increasing the recording duration from 3 to 30 seconds led to improved signal consistency, irrespective of the sensor’s connection status.

## Discussion

In this study, we present an original approach to acoustic volumetry that reimagines the simplest sensor configuration, fundamentally altering both the recording and computational methodologies. In addition, we developed dedicated, user-friendly freeware to facilitate its widespread application. To simplify the sensor, the microphone unit was omitted (comparable to Lao’s 1978 version); that is, instead of employing a dedicated detecting unit that converts the sound wave into a digital signal, only a single excitation unit —capable of propagating the sound wave within the measurement chamber— is retained, even when that unit incorporates a dynamic microphone cartridge with a highly sensitive, lightweight diaphragm. It is important to distinguish between the conventional concept of a “microphone” as an acoustic transducer and the functional role of the sensor’s diaphragm in this configuration.

The reimagined sensor, particularly in its circuit arrangement —where a single sound output is split into two paths, one driving the sensor and the other routed back to the sound card via the microphone input— demonstrated effective acoustic behavior suitable for volumetry. During characterization, the sinusoidal acoustic signal generated by the sound card’s headphone output was consistently recorded with a phase inversion (Fig. 3). This inversion results from the sensor’s diaphragm response; as the diaphragm moves forward in response to the acoustic stimulus, it increases the chamber pressure and produces a voltage of opposite polarity. Consequently, the recorded signal appears in anti-phase with respect to the original waveform, confirming the bidirectional acoustic-electrical conversion inherent in the sensor structure. Furthermore, independent of variations in transducer material or model, the sensor reliably detected a distinct resonance convexity (Fig. 4), providing additional evidence of its natural acoustic responsiveness.

In our sensor system, the lightweight and highly sensitive diaphragm of the acoustic transducer converts dynamic pressure variations within the chamber into an electrical signal. This relationship is described by the equation:

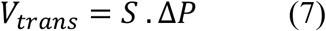

where *V*_*trans*_ represents the voltage generated by the transducer (sensor), *S* is the sensitivity (in volts per pascal) of the specific microphone capsule used, and Δ*P* denotes the acoustic pressure variation. The sensitivity *S* is a fixed characteristic of the specific microphone capsule used, and under stable operating conditions and within a defined frequency range, it is considered constant. Therefore, any variation in the measured output voltage *V*_*trans*_ reflects a corresponding change in the acoustic pressure inside the chamber —such as those caused by differences in internal volume or the presence of inserted sample objects that influence the chamber’s resonant behavior.

Since the sensor is connected in parallel with the microphone input of the sound card, the voltage across both branches is the same. According to Ohm’s law, the current flowing into the microphone input can be expressed as:

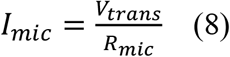

where *R*_*mic*_is the input resistance of the microphone port. Consequently, variations in acoustic pressure —through *V*_*trans*_ — result in corresponding changes in the input current *I*_*mic*_. This principle underpins the detection and quantification of internal sensor conditions, such as variable enclosed volume, through electrical measurements at the microphone input.

Although the direct or converted forms of the single recorded values of root mean square (RMS) amplitude (in decibels) have been used in previous studies (e.g. Kobata et al., 2004, and Sydoruk et al., 2020) for estimating sample volume, our pre-analyses (data not shown) indicated that, at least for the present sensor, predicting object volume based on shifts in the system resonance peak provides a more reliable and consistent approach. Observations (Fig. 5) showed that changes in the sampling object volume induce measurable shifts in the resonance peak. Accordingly, we developed a new computational method in which the resonance peak is determined using various mathematical models. Among these, the simple quadratic and Lorentzian models effectively estimated the peak, yielding more precise volumetry when modeling the overall trend of peak changes with different calibration sample sizes. The overall trend exhibited a robust linear relationship, with RMSE values of 1.980 μL using a linear fit and 1.662 μL for a logarithmic fit, thereby facilitating straightforward volume estimation.

It should be noted that our approach was designed to leverage inexpensive, readily available equipment; thus, while the current precision levels are adequate for many applications, there is promising potential for further improvement. Such precision may be further enhanced by employing higher-quality transducers (e.g., professional dynamic microphone cartridges) and sound cards (see Fig. 9), or even manufacturing exclusively designed circuits and transducers. Moreover, increasing the measurement time inherently increases the number of recorded acoustic cycles—serving as experimental replicates—and thus can statistically reduce measurement error; for instance, each additional second of playing and recording at 800 Hz provides an extra 800 sinusoidal waves for analysis (also see Fig. 9C). The current precision levels of the introduced sensor and platform are well-suited for applications in phenotyping, crop physiology, and other fields where multi-object sampling is common. For example, when 10 wheat grains were measured (Fig. 8), the total error (1.980 or 1.662 μL) was distributed among the grains, resulting in an average error of less than 0.2 μL per grain —a level of precision that is adequate for most physiological or phenotyping analyses. Moreover, reliable volumetry can be achieved with as few as three distinct frequency measurements, requiring a total recording duration of only 15 seconds (Fig. 8). Finally, additional technical considerations —such as reducing the sensor chamber’s capacity to better match the size of the sample and minimizing the air volume— may further enhance precision. For guidelines on measurement standards and to improve accuracy, refer to Sydoruk et al. (2020), where it is recommended that the working frequency should not exceed a wavelength less than four times the largest distance within the chamber to avoid uncertainties caused by acoustic effects such as standing waves and diffraction. Overall, the novel sensor and volumetry platform offer researchers across various scientific disciplines a reliable, fast, easy-to-use, and cost-effective tool for volumetric analysis.

## Conclusion

This study introduces a groundbreaking DIY acoustic volumetry platform that departs from traditional configurations by reviving microphone-free sensors. Our approach not only simplifies both the sensor and circuit architectures by harnessing the inherent dual acoustic behavior of dynamic microphone cartridges, but also provides an effective computational method for achieving precise and rapid volumetric measurements. By capitalizing on the shift in resonance peaks within a sealed chamber, the platform reliably estimates sample volumes with accuracy sufficient for applications in crop physiology and high-throughput phenotyping.

The integration of a streamlined circuit arrangement and a custom-developed Python-based freeware underscores the system’s accessibility and ease of adoption for researchers worldwide. Despite leveraging low-cost, commercially available components, the device demonstrates robust performance, as evidenced by a strong linear correlation in calibration and minimized measurement errors even with short acquisition times. These characteristics position the platform as a promising tool for volumetric analyses where conventional methods may be impractical or prohibitively complex.

Moreover, the device’s modular design opens avenues for further optimization. Potential enhancements could include employing higher-grade transducers and tailored circuitry to refine measurement precision, thereby broadening its applicability across diverse scientific disciplines. Future work may focus on adapting the platform to a range of sample types and volumes, reinforcing its utility in global phenotyping and beyond.

In summary, our microphone-free acoustic volumetry method offers a compelling balance of simplicity, cost-effectiveness, and accuracy. It represents a significant advancement in quantitative measurement techniques, providing a versatile tool that can be readily integrated into various research workflows, and setting the stage for further innovation in the field of digital volumetry.

## Acknowledgement

This study and the development of the Acoustic Volumetry freeware were carried out as part of the Easy Phenotyping Lab (EPL), a non-profit scientific initiative for sharing simple, reliable, and accessible plant phenotyping tools. For more information, see: https://haqueshenas.github.io/EPL. The English editing of the final manuscript and the code writing for the Acoustic Volumeter were both completed with assistance from ChatGPT-4o (OpenAI).

## Author Contributions

**A.H**.: Conceptualization, Methodology, Software, Formal Analysis, Writing.

**Y.E**.: Resources.

## Notes

https://github.com/haqueshenas/Acoustic-Volumeter

### Competing Interest Statement

The authors have declared no competing interest.

